# Multi-tissue patterning drives anterior morphogenesis of the *C. elegans* embryo

**DOI:** 10.1101/2020.04.27.064469

**Authors:** Stephanie Grimbert, Karina Mastronardi, Ryan Christensen, Christopher Law, David Fay, Alisa Piekny

**Affiliations:** Department of Biology, Concordia University, 7141 Sherbrooke Street West, Montreal, Quebec, Canada, H4B 1R6.; Laboratory of High Resolution Optical Imaging, NIH/NIBIB, 13 South Drive, Bethesda, Maryland, 20892.; Department of Molecular Biology, University of Wyoming, 1000 E. University Ave., Laramie, WY, 82071.

**Keywords:** *C. elegans*, morphogenesis, rosettes, polarity, cell migration, contractility, adhesion

## Abstract

Complex structures derived from multiple tissue types are challenging to study *in vivo*, and our knowledge of how cells from different tissues are coordinated is limited. Model organisms have proven invaluable for improving our understanding of how chemical and mechanical cues between cells from two different tissues can govern specific morphogenetic events. Here we used *Caenorhabditis elegans* as a model system to show how cells from three different tissues are coordinated to give rise to the anterior lumen. This poorly understood process has remained a black box for embryonic morphogenesis. Using various microscopy and software approaches, we describe the movements and patterns of epidermal cells, neuroblasts and pharyngeal cells that contribute to lumen formation. The anterior-most pharyngeal cells (arcade cells) may provide the first marker for the location of the future lumen and facilitate the patterning of the surrounding neuroblasts. These neuroblast patterns control the rate of migration of the anterior epidermal cells, whereas the epidermal cells ultimately reinforce and control the position of the future lumen, as they must join with the pharyngeal cells for their epithelialization. Our studies are the first to characterize anterior morphogenesis in *C. elegans* in detail and should lay the framework for identifying how these different patterns are controlled at the molecular level.

## Introduction

Our knowledge of how complex structures form in the developing embryo is limited due to challenges in studying the morphogenesis of multiple tissues simultaneously *in vivo*. *C. elegans* is an ideal organism to use for studies of development at the cellular and subcellular level, as they have multiple tissues with a defined number of cells. They are also a powerful genetic model, and the development of transgenic and microscopy tools makes them ideal for studying complex events *in vivo*. As the mechanisms regulating cell polarity and migration are highly conserved, studies of tissue morphogenesis in *C. elegans* have provided insight into how tissues develop in more complex organisms (Jacinto et al., 2001; Muller and Bossinger, 2003; Campanale et al., 2017).

Anterior morphogenesis is required for development of the anterior lumen, and involves the coordination of epidermal cells, pharyngeal cells and neuroblasts (neuronal precursor cells). Due to this complexity, it is not known how all three tissues form the anterior lumen, but revealing this could provide fundamental knowledge that is relevant for other complex developmental processes. The timing of anterior morphogenesis coincides with epidermal and pharyngeal morphogenesis, which are outlined below. The anterior-most epidermal cells migrate toward the anterior of the embryo after the epidermal cells meet at the ventral midline. Similarly, a large subset of pharyngeal cells polarize and form a cyst to define a lumen that aligns with the intestinal cells (Rasmussen et al., 2012). Anterior to the cyst are the arcade cells, which migrate anteriorly then move back into the embryo (Portereiko and Mango, 2001; Portereiko et al., 2004; Mango 2009). They also polarize and coordinate with the epidermal cells for epithelialization of the anterior pharynx, but it is not clear how this occurs (Von Stetina and Mango, 2015). During anterior morphogenesis, the neuroblasts presumably also undergo specific movements and patterning, but this has not been studied. Importantly, how all of these cell types are coordinated to give rise to a properly positioned anterior lumen remains a black box.

Epidermal morphogenesis has been relatively well-characterized, and requires chemical and mechanical signaling between different cell types. During mid-embryogenesis, the dorsal epidermal cells intercalate, followed by migration of the ventral epidermal cells toward the ventral midline to enclose the embryo through a process called ventral enclosure (Williams-Masson et al., 1997; Chisholm and Hardin, 2005). Ventral epidermal cell migration relies on precisely positioned neuroblasts for chemical and/or mechanical signaling (Bernadskaya et al., 2012; Ikegami et al., 2012; Wernike et al., 2016). EFN-VAB signaling is required for neuroblast positioning and for ventral enclosure, although it is not clear if the ligands and/or receptors are required in the neuroblasts or epidermal cells (George et al., 1998; Chin-Sang et al., 1999; Bernadskaya et al., 2012). Other signaling pathways implicated in controlling ventral epidermal cell migration include semaphorin (MAB-20) and plexin (PLX-2; Roy et al., 2000; Nakao et al., 2007; Ikegami et al., 2012). One model is that the receptors are expressed in the epidermal cells and receive cues from the neuroblasts or neighbouring epidermal cells to regulate branched F-actin assembly for their migration (Withee et al., 2004; Chisholm and Hardin, 2005; Patel et al., 2008; Patel and Soto, 2013; Wallace et al., 2018). In addition, subsets of neuroblasts in the middle-posterior of the embryo form rosettes, likely via planar cell polarity, to elongate the tissue in preparation for epidermal elongation. Rosettes are patterns required for the morphogenesis of different metazoan tissues, and form by apical-basal polarity or planar cell polarity, the latter of which are often associated with transient rosettes that facilitate tissue re-organization (Blankenship et al., 2006; Harding et al., 2014). The neuroblast rosettes that form in the ventral pocket may mechanically influence migration of the overlying epidermal cells during ventral enclosure (Wernike et al., 2016). Non-muscle myosin (NMY-2) is required in both the underlying neuroblasts and the epidermal cells for ventral enclosure, suggesting that these two tissues are mechanically connected (Wernike et al., 2016). Mechanical signaling also occurs between the epidermal and muscle cells to drive later stages of elongation, the subsequent step in epidermal morphogenesis (Gally et al., 2009). In a more recent study, the amphid neuronal precursors form lateral rosettes that maintain their organization as they migrate anteriorly with the epidermal cells, and there could be chemical or mechanical feedback through these cell-types (Fan et al., 2019). Neuroblast-epidermal cell interactions presumably also guide the anterior migration of epidermal cells for anterior morphogenesis.

Coincident with ventral epidermal morphogenesis, a large subset of pharyngeal cells polarize to form a cyst to define the pharyngeal lumen. These cells apically constrict with PAR-3 accumulated apically and laminin basally to form a cyst (rosette) with a central lumen (Rasmussen et al. 2012). Apical rosettes are formed by cells that apically constrict as a result of actomyosin activity, which is coordinated at the intercellular level via adhesion junctions (Sawyer et al., 2010; Martin and Goldstein, 2014). This gives rise to a bulb-like organization of cells with their narrow tips coming together to form a small hole apically (Harding et al., 2014). Apically formed rosettes are typically highly stable and ultimately give rise to lumens (Lecaudey et al., 2008; Nechiporuk and Raible, 2008). Interestingly, the anterior arcade cells remain distinct from the large pharyngeal cyst, yet must polarize to form a contiguous lumen (Portereiko and Mango, 2001; Portereiko et al., 2004; Mango 2009; Von Stetina and Mango, 2015). These cells must be precisely positioned in line with the more posterior pharyngeal cells and the anterior epidermal cells, but as indicated earlier, it is not clear how they do this.

Coordinated cell patterns, including rosettes, require that cells adopt polarity which is governed by intrinsic and extrinsic mechanisms. Apicobasal polarity occurs when complexes that include PAR-3, PAR-6 and PKC-3 localize apically, whereas other complexes such as those containing LET-413/Scribble are restricted basally (Etemad-Moghadam et al., 1995; Hung and Kemphues, 1999; McMahon et al., 2001). Many factors contribute to the onset and establishment of this asymmetric segregation of proteins, including the trafficking and turnover of active Cdc42, actomyosin organization, asymmetric distribution of specific phospholipids, adhesion junctions and mutual antagonism (Jiang et al., 2015; Campanale et al., 2017; Jewett and Prekeris, 2018; Motegi et al., 2020). In *C. elegans*, intestinal cells acquire apicobasal polarity, which is crucial for formation of the interior lumen (Leung et al., 1999; Bernadskaya et al., 201; Shafaq-Zadah et al., 2012; Rasmussen et al., 2012; Bossinger et al. 2015; Asan et al., 2016). Epidermal cells also have apicobasal polarity, which is reinforced by adhesion junction components that connect neighboring cells (Patel et al., 2008; Patel and Soto, 2013). These include the DAC (<u>D</u>LG-1 and <u>A</u>JM-1 <u>C</u>omplex) and the CCC (<u>C</u>atenin <u>C</u>adherin <u>C</u>omplex; E-cadherin [HMR-1], α-catenin [HMP-1] and β-catenin [HMP-2]) which are located sub-apically (Costa et al., 1998; Labouesse, 2006; Totong et al., 2007; Armenti and Nance, 2012; Pasti and Labouesse, 2015; Gillard et al., 2015; Sasidharan et al., 2018). Adhesion junctions connect actomyosin filaments intercellularly to coordinate cell movements and patterning. The CCC, but not the DAC, is also expressed in neuroblasts (Wernike et al., 2016) and the function of the CCC in these cells is not clear. The expression of different adhesion junction components in different cell types could influence adhesion and how mechanical forces are transmitted between cells.

Here we provide the first detailed description of *C. elegans* anterior morphogenesis, which involves the coordination of neuronal, epidermal and pharyngeal cells. Using diSPIM (dual-view inverted selective plane illumination microscopy) and confocal imaging, we observed that subsets of neuroblasts form concentric patterns around the site of the future lumen and that some of these neuroblasts ingress as the epidermal cells migrate anteriorly. We also observed the appearance of a PAR-6+, NMY-2+ and HMP-1+ focal point that marks the site of the future lumen and could represent apical projections from the pharyngeal arcade cells, which form a stable rosette aligned with the previously described pharyngeal cyst. Surrounding the central focal point, a subset of neuroblasts form specific patterns of PAR-6+, ANI-1+, NMY-2+ and HMP-1+ foci, and dorsal and ventral epidermal cell migration appears to be coordinated with these patterns. Removing subsets of neuroblasts affected the timing of lumen formation, whereas reducing the number of epidermal cells altered the position of the lumen. This in-depth description of anterior morphogenesis will provide a crucial starting point from which to explore the signaling pathways mediating these cell patterns and movements.

## Materials and Methods

### Strains

Strains were maintained on nematode growth medium (NGM) agar plates with a lawn of *Escherichia coli* (OP50) according to standard procedures (Brenner, 1974). The list of *C. elegans* strains used in this study is presented in Table 1. All strains were maintained at 20°C unless indicated otherwise. Genetic crosses were performed using standard protocols (for review, see Fay, 2005).

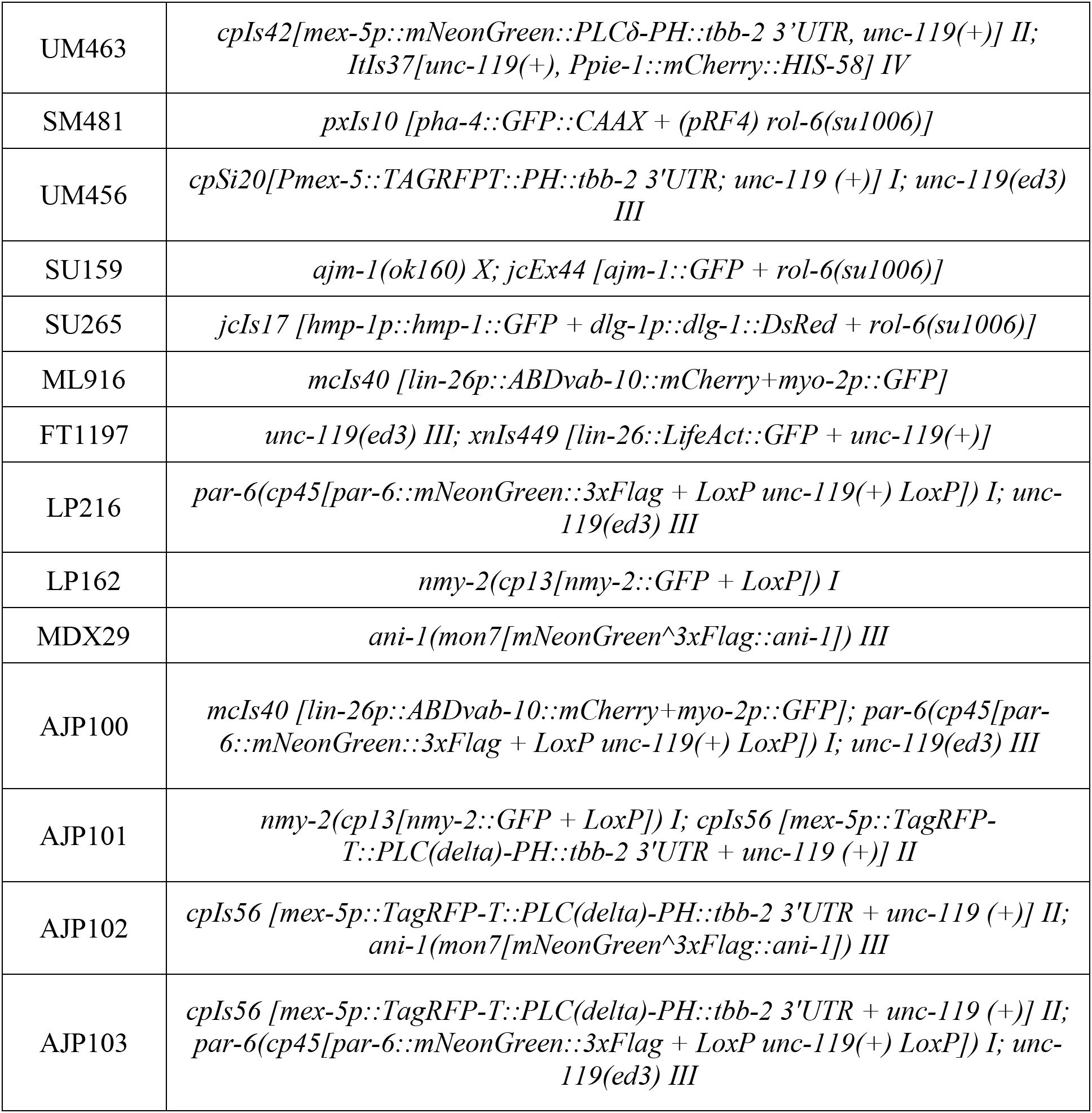
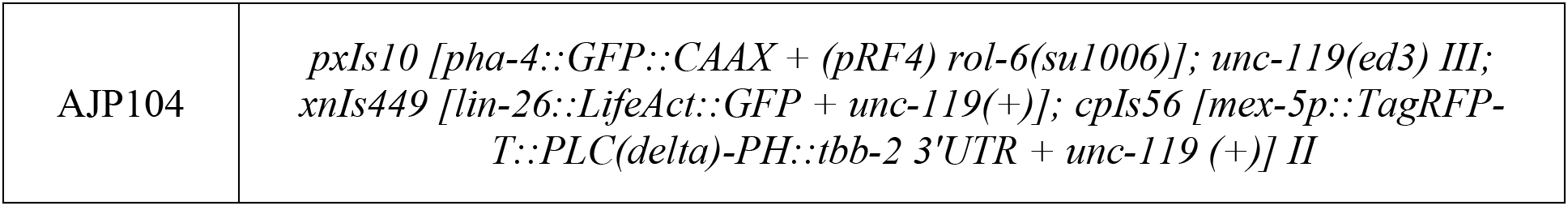

### RNAi

RNA-mediated interference (RNAi) by bacterial feeding was performed as described (Kamath et al., 2001; Timmons et al., 2001). Briefly, RNAi plates were made from NGM as above, with 50 mg/ml ampicillin and 1 mM IPTG. After growth to OD 0.6–1.0 (~6–12 hours), the cells were pelleted from their initial 5 mL volume and resuspended in varying amounts of LB to control the RNAi strength (*e.g.*, in 100–400 μL). Third and fourth larval stage (L3/L4) hermaphrodites were transferred to plates with *E. coli* (HT115) transformed with dsRNA constructs; the animals were incubated for 24 hours and then transferred to fresh RNAi plates, and progeny from these second plates were assessed for phenotype (Kamath et al., 2001). Y49E10.19 (*ani-1* RNAi) and W09C2.1 (*elt-1* RNAi) clones were used in this study (generously provided by J. C. Labbe, IRIC, Montreal and Michael Glotzer, University of Chicago, respectively).

### Microscopy

Imaging was performed on embryos collected as described (Sulston et al., 1983; Wernike et al., 2016). Images were acquired using a Nikon Eclipse Ti inverted microscope with a NIDAQ/Piezo stage, a 100× PlanApo lens (NA, 1.4), sweptfield confocal illumination (livescan; Bruker), an Ixon3 EMCCD camera (Andor) and NIS-Elements acquisition software. Fluorophores were excited using 488 nm and 561 nm lasers and a dual bandpass emission filter (520/20 nm + 630/30 nm). Z-stacks were collected every 0.5 μm, and embryos were imaged every 2 minutes for up to 2 hours. To limit phototoxicity and photobleaching, exposure times were kept below 300 milliseconds. For RNAi-treated embryos, Z-stacks were captured at 0.5 μm intervals every 4 minutes. For *en face* views, 0.2 μm Z-stacks were collected and acquired at intervals of 2–4 minutes. To image embryos *en face*, they were manipulated to be positioned vertically (upright) in small holes made in the agarose pads.

Images were also acquired using a fiber-coupled diSPIM (parts list and construction detailed in Kumar et al., 2014) with MicroManager software (Open-Source, Vale Lab UCSF). *C. elegans* embryos were mounted in an open-well chamber as described (Duncan et al., 2019). Image volumes were acquired in either single- or dual-view mode. Z-stacks were collected at 0.5 μm intervals with single-slice acquisition times of 12.75 milliseconds, leading to total volume acquisition times of 1.25 seconds for single-view volumes and 2.76 seconds for dual-view volumes. For single-view datasets, embryos were exposed simultaneously to 488 nm and 561 nm excitation with the emission optically split using a Hamamtsu W-View Gemini image splitter. Multi-point mode was used to capture multiple, spatially separated embryos per imaging session. For dual-view datasets, images were acquired sequentially, although within each path excitation at 488 nm and 561 nm was simultaneous and was again optically split with the Hamamtsu W-View Gemini image splitter. Dual-view data were also acquired using multi-point mode with multiple embryos captured per imaging run.

HILO, modified TIRF imaging, was performed using an inverted Nikon Ti-E microscope outfitted with a NI-DAQ piezo Z stage (National Instruments), an Evolve (EMCCD) camera, with Elements 4.0 acquisition software (Nikon), filters for 488 and 561 laser diodes, and a 100x CFI Apo TIRF objective. Images were exported as TIFFs and opened in Image J (NIH Image) to create Z-stack projections, perform image rotation and to crop desired regions.

### Image Analysis

Data files obtained by sweptfield confocal imaging were deconvolved using AutoQuant X3 (MediaCybernetics) with adaptive point-spread function (PSF) and blind deconvolution. The total number of iterations was set to 10 and the noise level was set to medium. All measurements were performed in FIJI (Fiji Is Just ImageJ; NIH) using the deconvolved images. The datasets were transferred to FIJI to generate hyperstacks and/or maximum-intensity projections and exported as TIFF files.

Single-view data obtained from diSPIM were exported as TIFF stacks through MicroManager software, and are shown in Fig. 1C. Dual-view data were deconvolved after export from MicroManager using custom fusion and deconvolution software (Guo et al., 2019). All datasets were then screened and subsequently analyzed.

**Figure 1.**
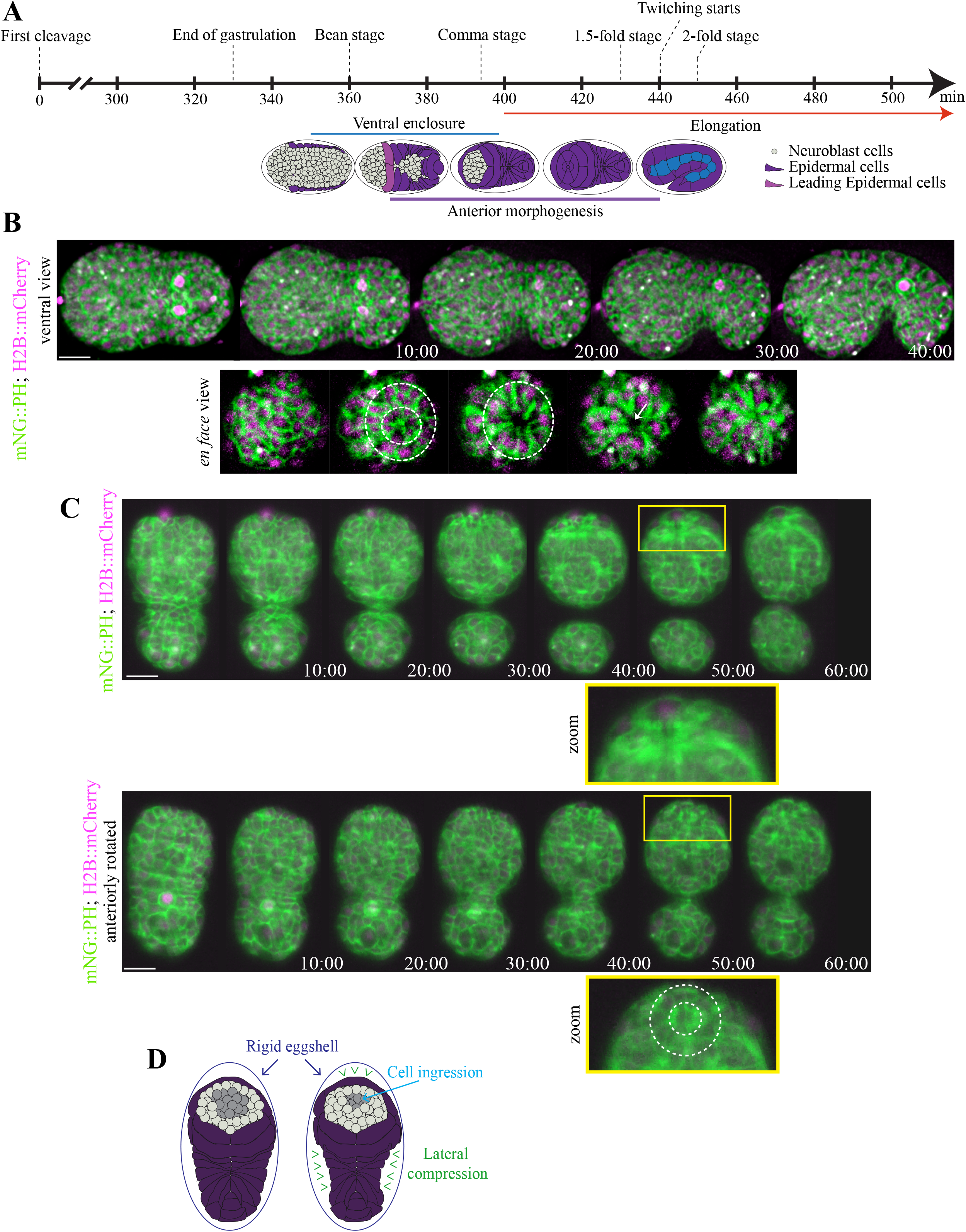
Cells ingress during anterior morphogenesis. **A)** The timeline (in minutes) shows the key morphogenetic events that occur during *C. elegans* embryogenesis. Anterior morphogenesis begins when the leading pair of migrating epidermal cells meet at the ventral midline (*t* = ~380), and continues throughout ventral enclosure and part of elongation (~440 minutes/twitching stage). **B)** Time-lapse images obtained by sweptfield microscopy (ventral view; anterior to the left) show an embryo co-expressing mNeonGreen∷PH to visualize membranes (green) and H2B∷mCherry to visualize nuclei (magenta) through anterior morphogenesis (time in minutes). The *en face* view shows cell organization at the anterior of the embryo, which results in the formation of a ‘ring in a ring’ (~430 minutes; white dashed circles). The star-like pattern in the center marks the site of the future lumen, which forms after cells ingress into the embryo. The scale bar is 10 μm. **C)** Raw time-lapse single-view images (ventral view; anterior pointing up) acquired using diSPIM show embryos as in B during anterior morphogenesis (time in minutes). The top panels show cell ingression at the anterior, whereas the bottom panels are tilted (anterior towards the reader) to show cell patterns (zoom, white dashed circles). The scale bar is 10 μm. Zoom images have been magnified by 200%. **D)** Cartoon schematics show the embryo inside the eggshell. The epidermal cells are shown in purple, the ingressing cells are shown in grey, and arrowheads (green) point to the forces generated laterally to cause ingression anteriorly.

The circumference of the arcade-cell rosette and cell length measurements were performed on Z-stack projections from 1.5-fold embryos expressing *pha-4p*∷GFP∷CAAX using FIJI as shown in Figure 2D. To measure the rosette diameter, *en-face* images were used, and three lines drawn across the diameter were measured for each embryo and averaged together. To measure cell length, lateral-view images were used, and the length of the most in-focus cell was measured from top to bottom. All measurements were converted from pixels to micrometers.

**Figure 2.**
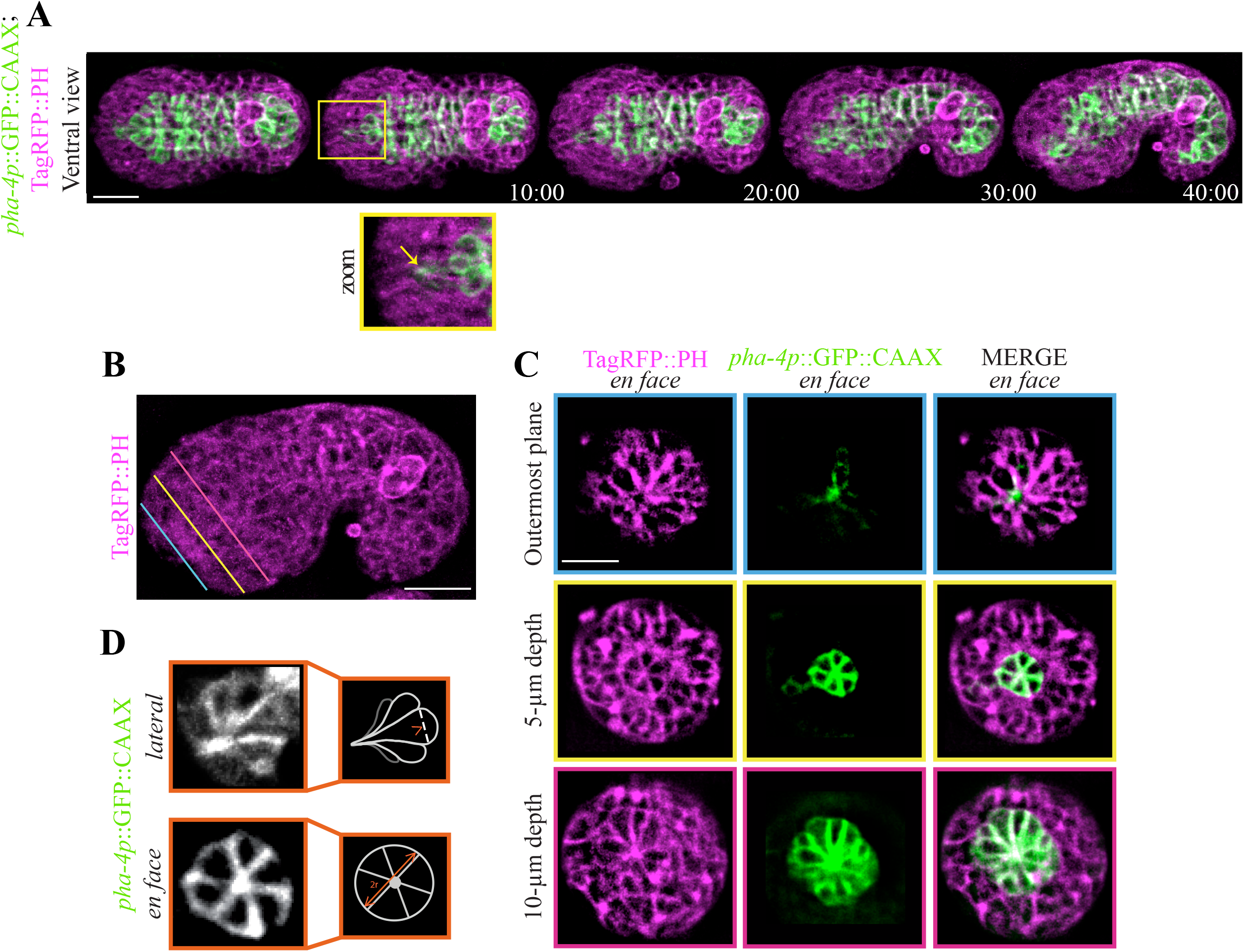
The pharyngeal arcade cells form a rosette. **A)** Time-lapse images acquired using sweptfield microscopy show an embryo co-expressing *pha-4p*∷GFP∷CAAX to visualize the pharyngeal cells in green and Tag-RFP∷PH to visualize membranes in magenta during anterior morphogenesis (time in minutes). The anterior-most pharyngeal cells (arcade cells) form apical constrictions (yellow box) which coalesce (zoom; yellow arrow). **B)** Images show an embryo ~ the 1.5-fold stage expressing Tag-RFP∷PH to visualize membranes (magenta). The colored lines indicate the three depths in the embryo corresponding to the images in C. **C)** Images show separate planes and merge from an embryo co-expressing Tag-RFP∷PH (magenta) and *pha-4p*∷GFP∷CAAX to visualize pharyngeal cells (green). Three *en face* views are shown, which correspond to the depths (colored lines) in B. The outermost plane (blue) shows the coalescence of the arcade-cell projections at the anterior of the embryo. At a 5 μm depth into the embryo (yellow) a small multicellular pharyngeal rosette is visible. The larger, previously characterized pharyngeal rosette is visible at a depth of 10 μm (pink). **D)** The regions used to determine the diameter of the anterior rosette and the length of individual cells are shown.

To determine the effects of *ani-1* RNAi on epidermal cell migration, we used NIS-Elements Viewer (Nikon) to measure the time needed for the amphid foci to reach their anterior location as a read-out for epidermal cell migration. The ‘start’ of anterior morphogenesis (*t* = 0 minutes) was when the leading pair of ventral epidermal cells met at the ventral midline (Figure 1A). Control and *ani-1* RNAi embryos were analyzed, and the phenotypes were categorized based on the time required for amphid cell migration (Figure 6). A chi-square test was used to determine statistical significance (*p* < 0.001) between control and *ani-1* RNAi phenotypes, as well as the proportion of phenotypes in each category with pharyngeal epithelialization (*p* < 0.01). Epithelialization was determined based on the accumulation of PAR-6 in the pharyngeal cells.

We determined the effects of *elt-1* RNAi on lumen position using deconvolved movie files that were analyzed with FIJI. Embryos were measured at *t* = 20 minutes after the leading pair of ventral epidermal cells met at the midline. The ratio of the distance between the Z2/Z3 cells (germline precursor cells) and the bright focal point (subset of PAR-6+ cells) and the total length of the embryo was determined for each embryo. Although there was no difference in the average ratio per se, an F-test revealed that the variability was significantly higher in the *elt-1* RNAi embryos (*p* < 0.01).

All figures were generated using Adobe Photoshop and Illustrator, after generating 8-bit format images using FIJI.

## Results

### Dynamic cell movements and patterning occur during anterior morphogenesis

We set out to characterize the cell movements and patterns that occur during anterior morphogenesis of *C. elegans* embryos, an event that has not been extensively studied. Anterior morphogenesis initiates during ventral enclosure when the leading pair of ventral epidermal cells meet at the ventral midline (Fig. 1A) and continues until the 1.7-fold stage of embryogenesis when the muscles start twitching (Chisholm and Hardin, 2005). Initially the anterior-most part of the embryo is composed primarily of neuroblasts, and the epidermal cells migrate anteriorly to cover the head in epidermal tissue (Chisholm and Hardin, 2005). To visualize the cell patterns and movements during anterior morphogenesis, we imaged embryos co-expressing mNeonGreen-tagged pleckstrin homology (PH) domain (mNG∷PH), which localizes to membranes and mCherry-tagged H2B (H2B∷mCherry) to visualize nuclei (Fig. 1B-D). We used two types of imaging methods—sweptfield confocal microscopy (Fig 1B; *n* = 37) and diSPIM (Fig. 1C; Movie S1; *n* = 8) to visualize changes in cell movements and patterns at different resolutions. diSPIM captures two orthogonal image volumes for two different views of the sample at each time point, allowing computational processing to create a single isotropic volume with better resolution than traditional imaging systems. Single-view images are shown here (Fig. 1C). At 10–15 minutes after the start of anterior morphogenesis, rings of neuroblasts formed around the site of the future lumen (Fig. 1B-C). As the epidermal cells migrated toward the anterior, a subset of these neuroblasts underwent ingression (Fig. 1B-C). To better visualize the cell movements occurring during anterior morphogenesis, we imaged embryos *en face* (Fig. 1B; see Methods). Using this approach, we were able to more clearly observe the ring-like patterns formed by the neuroblasts during anterior morphogenesis. The ingression of cells likely occurred due to forces generated by epidermal constriction, as the embryo begins to elongate and lengthen at this time (Fig. 1D; Chisholm and Hardin, 2005). In addition, a large proportion of neuroblasts will differentiate into neurons that form the nerve ring, which encircles the pharynx, and this inward movement likely helps position these neuronal precursors appropriately (Hobert, 2010). When the epidermal cells reached the anterior region, the lipids revealed a star-like pattern, likely corresponding to growing axons, with the central point corresponding to the site of the future lumen (Fig. 1B-C). Thus, several distinct cell movements occurred during anterior morphogenesis that require further characterization.

### The arcade cells of the pharynx form a stable rosette

While we were imaging embryos using the membrane marker, we observed a rosette that formed at a depth of ~ 4–5 μm and aligned with the site of the future lumen (Fig. 2A-C). This rosette formed anteriorly to the larger, previously described cyst formed by pharyngeal cells (Fig. 2C; Rasmussen et al. 2012). This rosette, which contains six cells, persisted for an extended period of time, which is characteristic of apical polarity–derived rosettes known to give rise to lumens (Fig. 2C; Sawyer et al., 2010; Harding et al., 2014; Martin and Goldstein, 2014). To determine if these cells are pharyngeal and to follow their patterning more specifically, we imaged embryos co-expressing *pha-4p*∷GFP∷CAAX (*n* = 18). This probe localizes to the membranes of pharyngeal cells, as *pha-4* encodes a forkhead box A (FOXA) transcription factor required for pharyngeal cell fate, and CAAX is post-translationally farnesylated (farnesyl is a lipid moiety; Horner et al., 1998; Kalb et al., 1998; Gaudet & Mango 2002; Roberts et al., 2008; Manolaridis et al., 2013). At the start of anterior morphogenesis, the most anterior *pha-4*+ve cells moved toward the anterior of the embryo but subsequently moved posteriorly within the embryo while retaining a narrow region of *pha-4*+ve signal that aligned with the center of the rosette (Fig. 2A, C). Approximately 40-50 minutes after the start of anterior morphogenesis, these cells adopted a teardrop shape (Fig. 2A, D). Imaging *pha-4p*∷GFP∷CAAX and TagRFP∷PH 1.5-fold embryos *en face* revealed the rosette more clearly. Consistent with what we had observed earlier, the rosette was positioned at a depth of 5 ± 0.40 μm from the anterior tip of the embryo (Fig. 2C, middle panel), whereas the previously characterized, larger pharyngeal rosette was at 11 ± 0.40 μm (*n* = 13 embryos; Portereiko & Mango, 2001; Fig. 2C, bottom panel). To better characterize the small *pha-4*+ve rosette, we measured its diameter along three axes per rosette, as well as the length of individual cells in the rosette (Fig. 2D). We found that the average diameter was 6.9 ± 0.13 μm (*n* = 13 embryos), whereas the average cell length was 4.9 ± 0.04 μm (*n* = 16 cells), consistent with what we had observed for depth. Given the location and number of these cells, the anterior rosette is likely composed of arcade cells. These cells formed apical constrictions with projections that extended anteriorly and remained in place as the arcade cells moved back inside the embryo. As these projections were aligned with the center of the rosette, this is one of the first anterior-positioned markers that defines the site of the future lumen.

### Distinct patterns of foci form in the anterior region of the embryo

Next, we characterized patterns formed by the anterior epidermal cells and neuroblasts. We speculated that the neuroblasts form distinct patterns that could influence epidermal cell migration. We previously showed that the CCC component HMP-1/α-catenin is enriched during ventral enclosure in the mid-posterior neuroblasts that forms transient rosettes for tissue re-organization (Wernike et al., 2016). Imaging analysis of embryos expressing GFP-tagged HMP-1/α-catenin during anterior morphogenesis showed its enrichment in subsets of anterior neuroblasts, or other cells that form distinct patterns (Fig. 3A; *n* = 8). From 10–30 minutes, we observed the formation of two pentagons of HMP-1+ve foci on either side of a larger, bright focal point (BFP). The BFP could correspond to the apical projections of the arcade cells, and/or the adhesion of these projections to the nearby neuroblasts (Fig. 3A). Co-imaging of HMP-1∷GFP and DLG-1∷RFP, a DAC component that is part of the adhesion junctions and is expressed predominantly in epidermal cells (Costa et al., 1998), revealed that the anterior-most epidermal cells migrated toward the HMP-1-rich foci (Fig. 3A; *n* = 8). We also imaged embryos expressing AJM-1∷GFP, the other DAC component that is enriched in epidermal cells (Costa et al., 1998), which showed localization patterns similar to DLG-1 (Fig. 3A; *n* = 15). However, as this probe is brighter than DLG-1, we used Fire LUTs to better visualize changes in AJM-1 levels to reveal more detailed localization patterns. Fire LUTs convert the signal into different colors based on intensity levels, where white or red is high, and blue or violet is low. As anterior morphogenesis progressed, AJM-1 foci appeared at the ‘front’ of the anterior ventral epidermal cells, which could reflect the formation of adhesions with the underlying neuroblasts. Other more centrally located foci were initially faint but became brighter as the epidermal cells reached the anterior and coalesced. This likely reflects the maturation of junctions and/or sites of adhesion as cells make contacts with neighboring epidermal cells and/or neuroblasts or pharyngeal cells. Ultimately, the epidermal cells arranged themselves to form two bilateral rings, which fused and joined the pharynx, with another ring forming around it (Fig. S1A).

**Figure 3.**
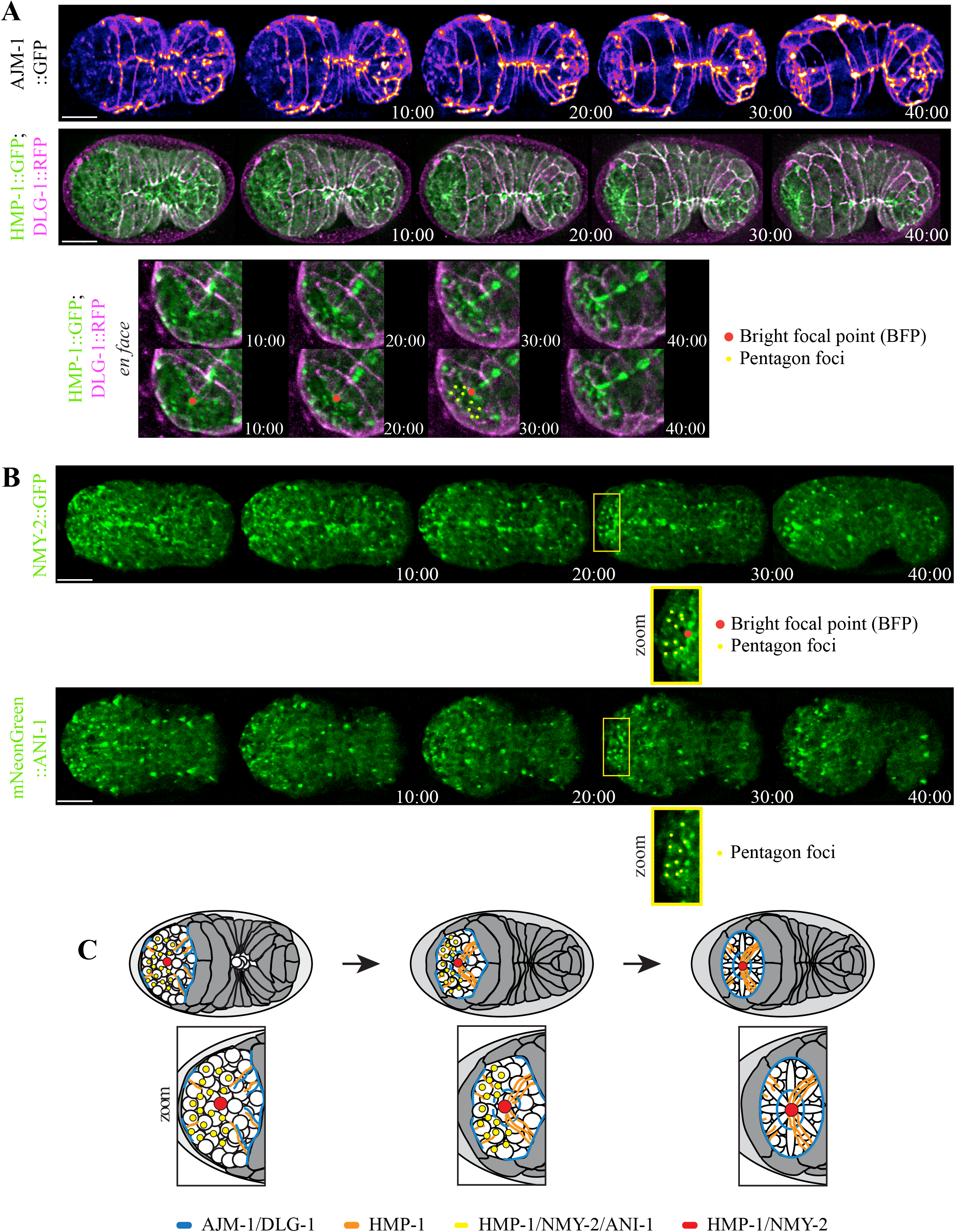
Adhesion junctions and contractility components are enriched in subsets of cells during anterior morphogenesis. **A)** Time-lapse images acquired using sweptfield microscopy show ventral views (anterior to the left) of embryos expressing AJM-1∷GFP (Fire LUTs; top panel) or HMP-1∷GFP (green); DLG-1∷RFP (magenta; middle panel) during anterior morphogenesis (time in minutes). For the Fire LUTs, cool colors, such as blue or violet correspond to lower levels of signal vs. warmer colors such as yellow or white, which show higher levels. The scale bar is 10 μm. The anterior region is shown more clearly in the bottom group of *en face* images. AJM-1 and DLG-1 demarcate epidermal cell boundaries, as well as sites where the migrating anterior epidermal cells contact the substrate. Signals associated with the anterior epidermal cells strengthen as they meet and form junctions in a circular pattern. In addition to the epidermal cells, HMP-1 (α-catenin) accumulates in subsets of neuroblasts and/or anteriorly positioned pharyngeal cells in a distinct pattern—a centrally located BFP (*en face*; red circle) and two bilaterally positioned pentagons on either side of this brighter signal (*en face*; yellow circles). **B)** Time-lapse images acquired using sweptfield microscopy show ventral views (anterior to the left) of embryos expressing NMY-2∷GFP (top panel) or mNeonGreen∷ANI-1 (bottom panel). The scale bar is 10 μm. Zoomed-in regions (yellow box; 150%) show the patterns in the anterior region more clearly. **C)** Cartoon schematics show patterns of the DAC and CCC junction components during anterior morphogenesis, as well as NMY-2 and ANI-1. AJM-1 and DLG-1 are in blue, HMP-1 is in orange, the pentagon foci are in yellow (HMP-1, NMY-2 and ANI-1) and the bright focal point (BFP) is in red (HMP-1 and NMY-2).

As mentioned above, the enrichment of HMP-1 within subsets of cells could reflect their adhesion with neighboring cells or could indicate that these cells have polarity. As actomyosin is typically enriched at junctions, we imaged embryos expressing GFP∷NMY-2 (non-muscle myosin; Fig. 3B; *n* = 23). We also imaged GFP∷ANI-1 (anillin), which we previously found to be specifically enriched in neuroblasts (Wernike et al., 2016), and is an indicator of polarity during cell division as it binds to actomyosin (Fig. 3B; *n* = 29). Indeed, both NMY-2 and ANI-1 localized to the pentagon foci, supporting the hypothesis that these foci are associated with neuroblasts (Fig. 3B, C). However, NMY-2, but not ANI-1, localized to the BFP that was enriched with HMP-1. This signal could reflect the apical projections emanating from the arcade cells or unique enrichments at sites of contact with neuroblasts that are not ANI-1+ve (Fig. 3B, C).

We also observed a semi-circle-like pattern of foci that arise ventrally to the BFP and that align with the leading edge of the anterior ventral epidermal cells (Fig. 3A; better visualized in 5A). The majority of markers localized to these foci with the exception of ANI-1. Although it is not clear if these foci correspond to epidermal cells or neuroblasts, they likely reflect contact points between the two cell types. Some proteins such as AJM-1 are more strongly enriched in epidermal cells vs. HMP-1, which may be expressed in both cell types. It is not clear why ANI-1 is not at these sites. As an organizer of actomyosin filaments, ANI-1 could help control polarity but may not have a role in cell-cell adhesion in these cell types.

### Ventral and dorsal epidermal cells migrate to an anterior-ventral position

Next, we characterized anterior epidermal cell migration. To do this, we monitored the localization of F-actin expressed exclusively in epidermal cells during anterior morphogenesis in embryos expressing either mCherry-tagged VAB-10 actin-binding domain (ABD; *n* = 32) or GFP-tagged LifeAct (*n* = 12), each of which was driven by an epidermal promoter (*lin-26;* Gally et al., 2009; Zilberman et al., 2017). As the anterior ventral epidermal cells migrated more anteriorly, we observed the formation of actin-rich projections (Fig. 4A, zoom). As the cells reached their anterior-most position and the embryos began to elongate (20–30 minutes), the actin projections pointed toward a more central, ventrally-shifted location coincident with the site of the future lumen (Fig. 4A, zoom). The dorsal epidermal cell also migrated during this time, moving over a greater distance to the site of the more ventrally positioned lumen (Fig. 4A, zoom).

**Figure 4.**
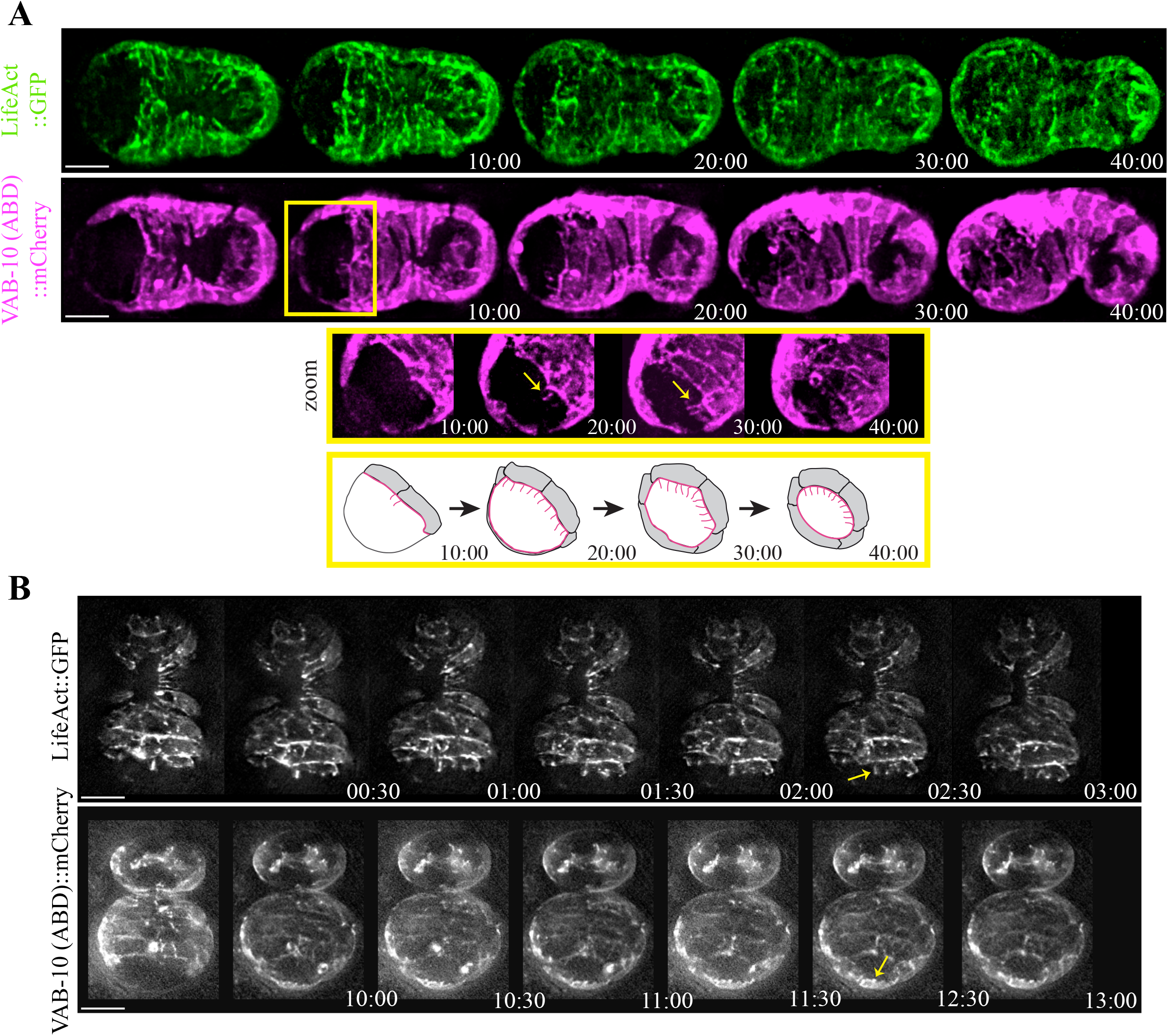
Projections form at the leading edge of the migrating anterior ventral epidermal cells. **A)** Time-lapse images acquired using sweptfield microscopy show ventral views (anterior to the left) of embryos expressing *lin-26p*;LifeAct∷GFP (green) or *lin-26p*;VAB-10 (actin-binding domain; ABD)∷mCherry (magenta) during anterior morphogenesis (time in minutes). The scale bar is 10 μm. The ABD from VAB-10 and LifeAct both indicate F-actin and are expressed exclusively in epidermal cells using the *lin-26* promoter. At the onset of anterior morphogenesis, the anterior of the embryo is devoid of epidermal cells, which subsequently migrate into this space. The ventral cells migrate a shorter distance as compared with the dorsal cells. The yellow box highlights a region that has been tilted to better visualize the F-actin projections (yellow arrows). The bottom row shows cartoon schematics that indicate the orientation of the projections in pink. **B)** Time-lapse images acquired by HILO microscopy show ventral views (anterior pointing down) of embryos expressing *lin-26p*;LifeAct∷GFP (top panel) or *lin-26p*;VAB-10(ABD)∷mCherry (bottom panel; time in minutes). The scale bar is 10 μm. The projections are dynamic and longer at the leading edge of the ventral epidermal cells (top panel; white arrow), whereas the dorsal epidermal cells have smaller, more uniform projections (bottom panel; white arrow).

To follow epidermal cell migration with a higher temporal resolution, we imaged F-actin in embryos using HILO (highly inclined and laminated optical sheet) microscopy (Wernike et al., 2016). With this method, TIRF (total internal reflection fluorescence) objectives are used and the illumination beam is angled to obtain a thicker *z* plane (*e.g.*, in this case to a depth of 1-2 μm) emerging obliquely from the sample. In Fig. 4B, the top panel shows images of LifeAct taken every 30 seconds (anterior is pointing down; Movie S2; *n* = 8). Projections from the anterior ventral epidermal cells were dynamic, extending anteriorly for several microns. However, they did not extend far past the leading edge of the cell. The bottom panel shows images of an embryo expressing VAB-10 (ABD)∷mCherry that were taken every 30 seconds, in which the dorsal epidermal cells were more visible (anterior pointing down; Movie S3; *n* = 9). The extensions at the leading edge of the most anterior dorsal cell were more uniform and smaller in length as the cell moved toward the site of the future lumen as compared with the ventral epidermal cells (Fig. 4B). It is striking how both subsets of epidermal cells coordinated their movements to reach the ventrally positioned future lumen at the same time, suggesting that their movements could be controlled by cues associated with other cells in this location.

### PAR-6+ve foci are coupled with the migration of epidermal cells

As subsets of neuroblasts formed distinct patterns during anterior morphogenesis, we next determined how these patterns correlate with anterior epidermal cell migration. To do this, we imaged embryos expressing mNeonGreen∷PAR-6, which localizes apically, and mCherry-tagged ABD from VAB-10 to visualize epidermal F-actin (*n* = 15; Wernike et al., 2016; Fan et al., 2019). As reported recently by the Bao lab, we observed that PAR-6 was enriched at the vertex of the amphid dendrite tips (Fig. 5A; Fan et al., 2019). While their images also revealed additional foci enriched in the anterior, these were not characterized. We saw PAR-6 localize to the BFP, the pentagon foci and in the semi-circle of foci (Fig. 5A). In particular, the PAR-6+ve BFP appeared on the ventral side of the embryo ~10 minutes after the start of anterior morphogenesis and at the same location where we had observed HMP-1 (Fig. 3A, 5A). We propose that the BFP corresponds to the apical projections of the arcade cells. In support of this, a PAR-6+ve BFP was no longer visible in embryos treated with *pha-4* RNAi to disrupt pharyngeal cell fate (Fig. S1B; Kalb et al. 1998). By ~20 minutes after the start of anterior morphogenesis, we saw a PAR-6+ve semi-circle of foci corresponding to the leading edge of the migrating epidermal cells (Fig. 4A, zoom). The projections of the epidermal cells did not cross the BFP or semi-circle of foci. We also saw the two pentagon patterns that were initiated at a more dorsal position and then moved more ventrally to meet with the semi-circle (Fig. 5A). By 40 minutes, these foci had formed a rectangular pattern with the semi-circle and the three ventral foci from each pentagon marking one side and the two dorsal foci marking the other, which quickly resolved into a circular pattern around the lumen (Fig. 5A, zoom). We propose that subsets of neuroblasts signal and/or act as a substrate to direct the migration of the anterior epidermal cells. Their patterns could ensure that the cues are properly positioned so that the ventral epidermal cells migrate a shorter distance and/or more slowly as compared with the dorsal epidermal cells to properly position the lumen. The pharyngeal cells contributing to the bright focal point also could provide important cues to control epidermal cell migration.

**Figure 5.**
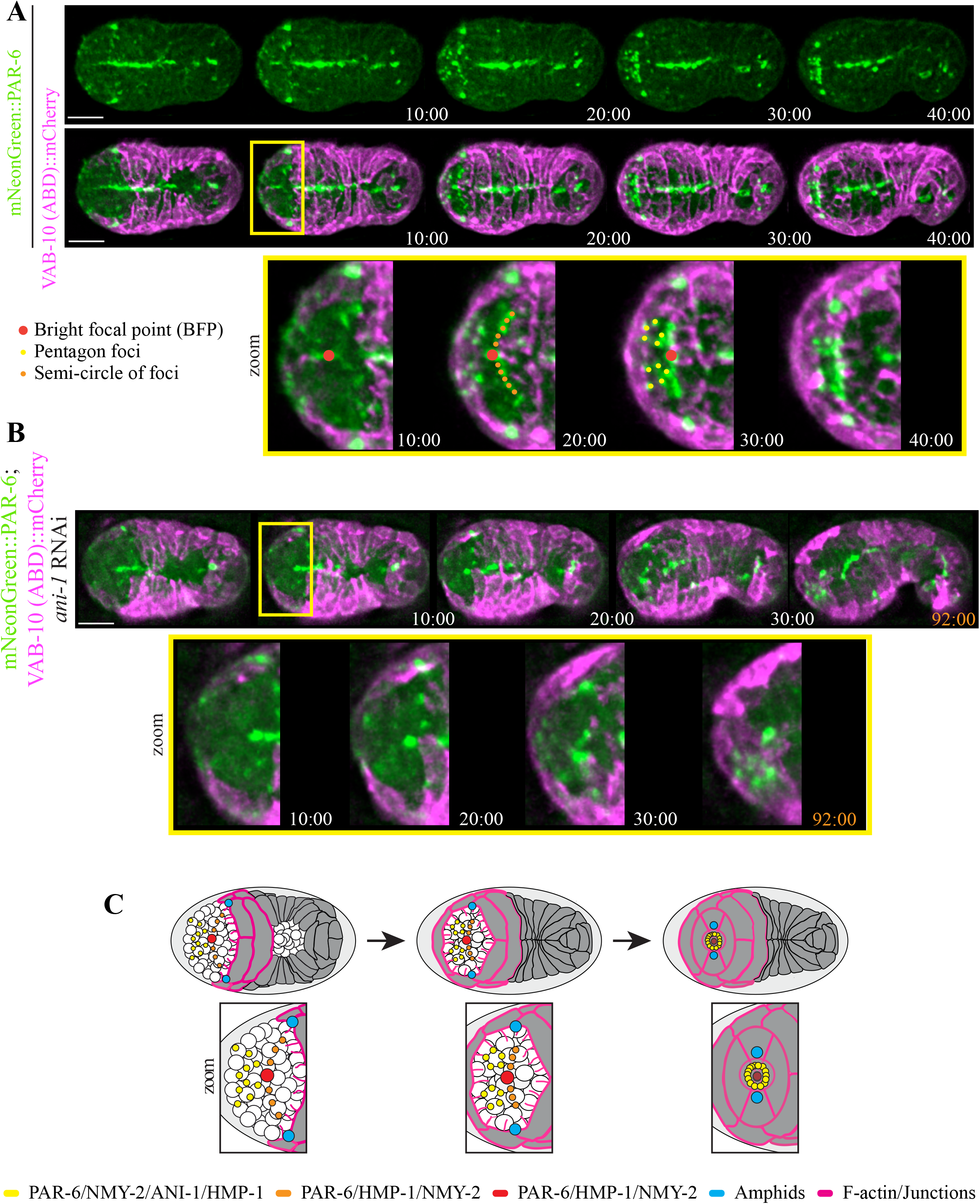
Distinct patterns formed by PAR-6+ cells correlate with the migrating anterior epidermal cells. **A)** Time-lapse images acquired using sweptfield microscopy show ventral views of an embryo expressing mNeonGreen∷PAR-6 (green; top panel) and *lin-26p*;VAB-10 (ABD)∷mCherry (magenta; middle panel) during anterior morphogenesis (time in minutes). The scale bar is 10 μm. Zoomed-in regions (bottom panel; 200%) show the patterns formed by PAR-6+ cells more clearly (yellow box). PAR-6 localizes to the BFP, which demarcates the site of the future lumen (zoom; red), the bilateral pentagon foci (zoom; yellow), and the semi-circle of foci that corresponds to the anterior-most boundary of ventral epidermal cells (zoom; orange). **B)** Time-lapse images acquired using sweptfield microscopy show ventral views of an embryo as in A after treatment with *ani-1* RNAi during anterior morphogenesis (time in minutes). Whereas the BFP is still present after *ani-1* depletion, some of the foci associated with the pentagons and the semi-circle of foci not visible. The scale bar is 10 μm. **C)** Cartoon schematics show the patterns of foci and F-actin during anterior morphogenesis as in Fig. 3 but also include the location of foci from PAR-6+ve cells during anterior morphogenesis (amphids are indicated in blue).

### Anterior epidermal cell migration depends on subsets of anterior neuroblasts

Next, we determined whether perturbing these localization patterns by reducing the number of neuroblasts impacts epidermal cell migration or lumen position. We previously showed that *ani-1* is required for neuroblast cell division (Fotopoulos et al., 2013; Wernike et al., 2016). Depleting *ani-1* in embryos co-expressing mNeonGreen∷PAR-6 and *lin-26p*;VAB-10 (ABD)∷mCherry caused changes in PAR-6 localization and delayed anterior epidermal cell migration (*n* = 34; Fig. 5B). We observed a correlation in the severity of anterior morphogenesis phenotypes with higher degrees of perturbed PAR-6 localization and delayed migration, including failed polarization of the anterior pharynx (Fig. 5B).

We quantified the different anterior morphogenesis phenotypes caused by *ani-1* RNAi in embryos co-expressing mNeonGreen-tagged PAR-6 and a Tag-RFP membrane marker (*n* = 15 control and *n* = 28 *ani-1* RNAi embryos; PH domain from PLCγ; Fig. 6A). To measure the duration of anterior epidermal cell migration, we determined the time it took for the amphid neurons to reach the anterior region. The Bao lab recently showed that these PAR-6+ve cells co-migrate with epidermal cells (Fan et al., 2019). We observed embryos with a range in delay phenotypes that we categorized as mild (50–69 minutes; 45%), moderate (70–89 minutes; 30%) and severe (≥90 minutes; 10%), based on the amphid migration time relative to that of control embryos (40–49 minutes; 15%; Fig. 6B). We also observed that PAR-6 failed to localize in the anterior pharynx in a subset of the *ani-1* RNAi embryos. Of the embryos that displayed a mild delay phenotype, 33% failed to polarize, whereas 67% of embryos with moderate delays failed and 100% of the severely delayed embryos failed (Fig. 6B). We correlated the changes in neuroblast/foci patterns with the observed phenotypes and found that more of the foci that form the semi-circle or pentagons were lost or severely disorganized as the delay worsened from mild to severe (Fig. 6C). It is important to note that we did not observe changes in the position of lumen formation (*e.g.*, in embryos with a less-severe delay) in *ani-1* RNAi embryos, nor was the BFP altered in intensity or appearance. This suggests that subsets of neuroblasts are required for controlling the speed of epidermal cell migration, but not necessarily their direction. Also, the failure to polarize the pharynx in the more severely delayed embryos likely reflects a threshold requirement for the timing of cell-cell contacts to complete lumen formation.

**Figure 6.**
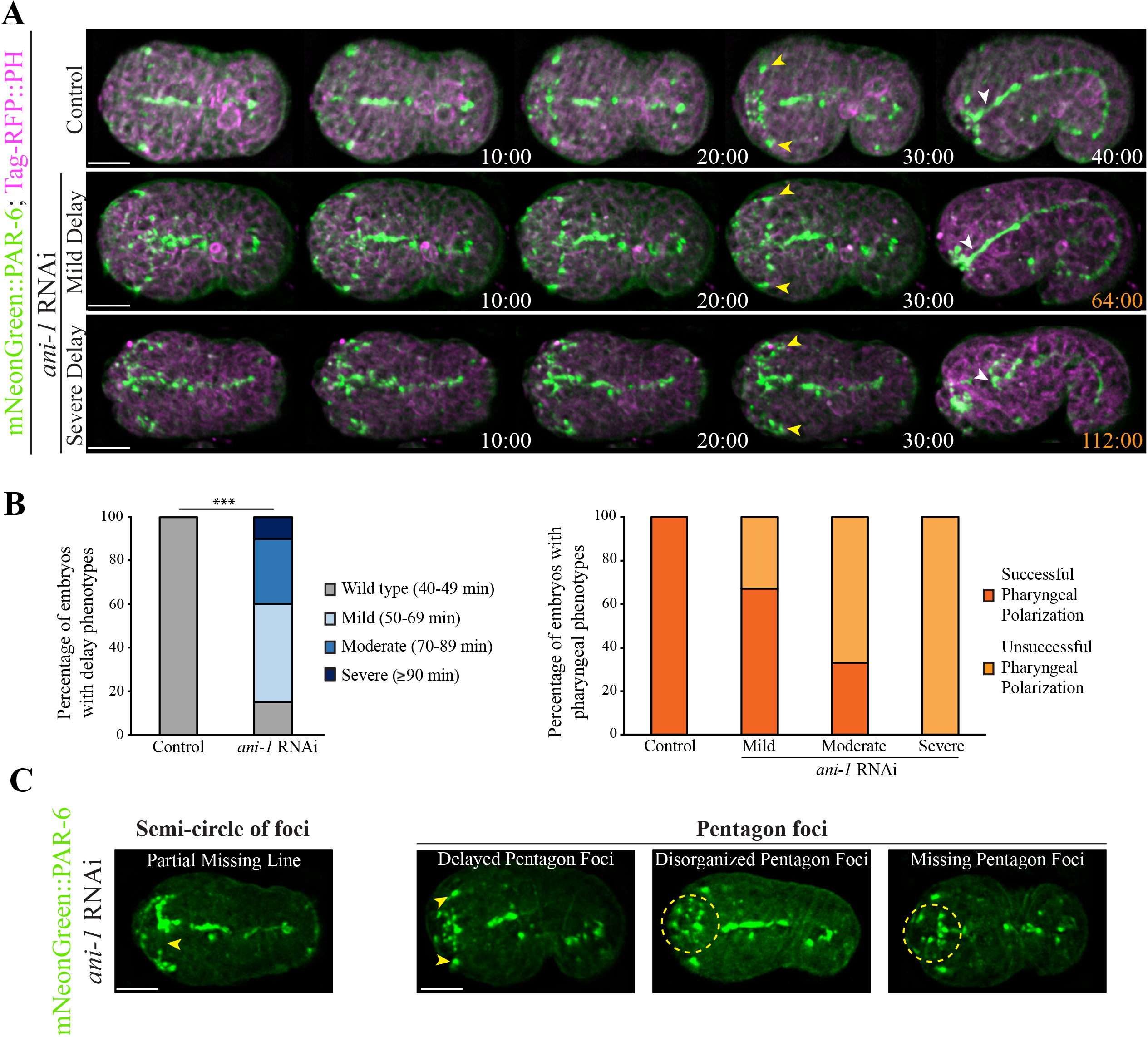
Disrupting neuroblast division causes delays in anterior epidermal cell migration. **A)** Time-lapse images acquired using sweptfield microscopy show ventral views of embryos expressing mNeonGreen∷PAR-6 (green); Tag-RFP∷PH (magenta) in control (top panel) and after *ani-1* RNAi to disrupt neuroblast cell division (middle and bottom panels; time in minutes). Delays in epidermal cell migration, marked by the amphids (yellow arrowheads), correlated with changes in the number or position of PAR-6+ve foci and/or failure of the anterior pharynx to polarize (white arrowheads) in *ani-1–*depleted embryos compared to control embryos. **B)** Epidermal cell migration was monitored by measuring the time it took for the amphids to reach the anterior region. The bar graph on the left shows the proportion of embryos with different delays in epidermal cell migration after *ani-1* RNAi when compared with control embryos. The bar graph on the right shows the proportion of control and *ani-1* RNAi embryos with delay phenotypes in which the anterior pharynx failed to polarize. **C)** Images show changes in the patterns of PAR-6+ve foci observed in mNeonGreen∷PAR-6 (green) embryos with epidermal cell migration delays after *ani-1* RNAi. The scale bar for all embryos is 10 μm.

### Epidermal cells determine the position of the lumen

Next, we determined how mild perturbation of epidermal cell fate affects anterior morphogenesis. To accomplish this, we partially depleted *elt-1* in embryos co-expressing mNeonGreen∷PAR-6 and Tag-RFP∷PH (*n* = 15 control vs. *n* = 35 *elt-1* RNAi). *elt-1* encodes a GATA-like transcription factor and is essential for determining epidermal cell fate (Fig. 7A; Page et al. 1997). After partial *elt-1* depletion, there was no significant delay in the migration of anterior epidermal cells. However, we observed a ventral shift in the position of the anterior lumen formation (Fig. 7 A, B). To quantify this, we measured the ratio of the distance between the lumen (marked by PAR-6) and the Z2/Z3 cells (germline precursors) to the total length of the embryo. Whereas the average ratio was not significantly different, the variability was significantly higher in mildly perturbed *elt-1* RNAi embryos (*n* = 19) as compared with control embryos (*n* = 10; Fig. 7B). In a subset of the more severely affected embryos we saw that the PAR-6+ve foci formed patterns that were initially similar to those of control embryos, but these foci failed to coalesce and shifted more ventrally as compared to control embryos (Fig. 7A). The BFP also underwent a more ventral shift in severely affected embryos, and the pharynx failed to epithelialize. Therefore, the position of the anterior lumen was determined by the epidermal cells. Although it is not clear why the lumen tended to shift more ventrally, this could be because there are two rows of dorsal cells that intercalate and fuse, and perhaps the loss of one/more of these cells has less of an impact vs. the loss of ventrally positioned cells. These cells also migrate over a longer distance and may interpret cues differently as compared with the ventrally positioned cells. It is interesting that subsets of neuroblasts also relied on epidermal cells for their positioning (Fig. 7A) suggesting that these tissues are interconnected. This is supported by findings from the Bao lab, which showed that the amphid neurons are connected with the migrating epidermal cells (Fan et al., 2019).

**Figure 7.**
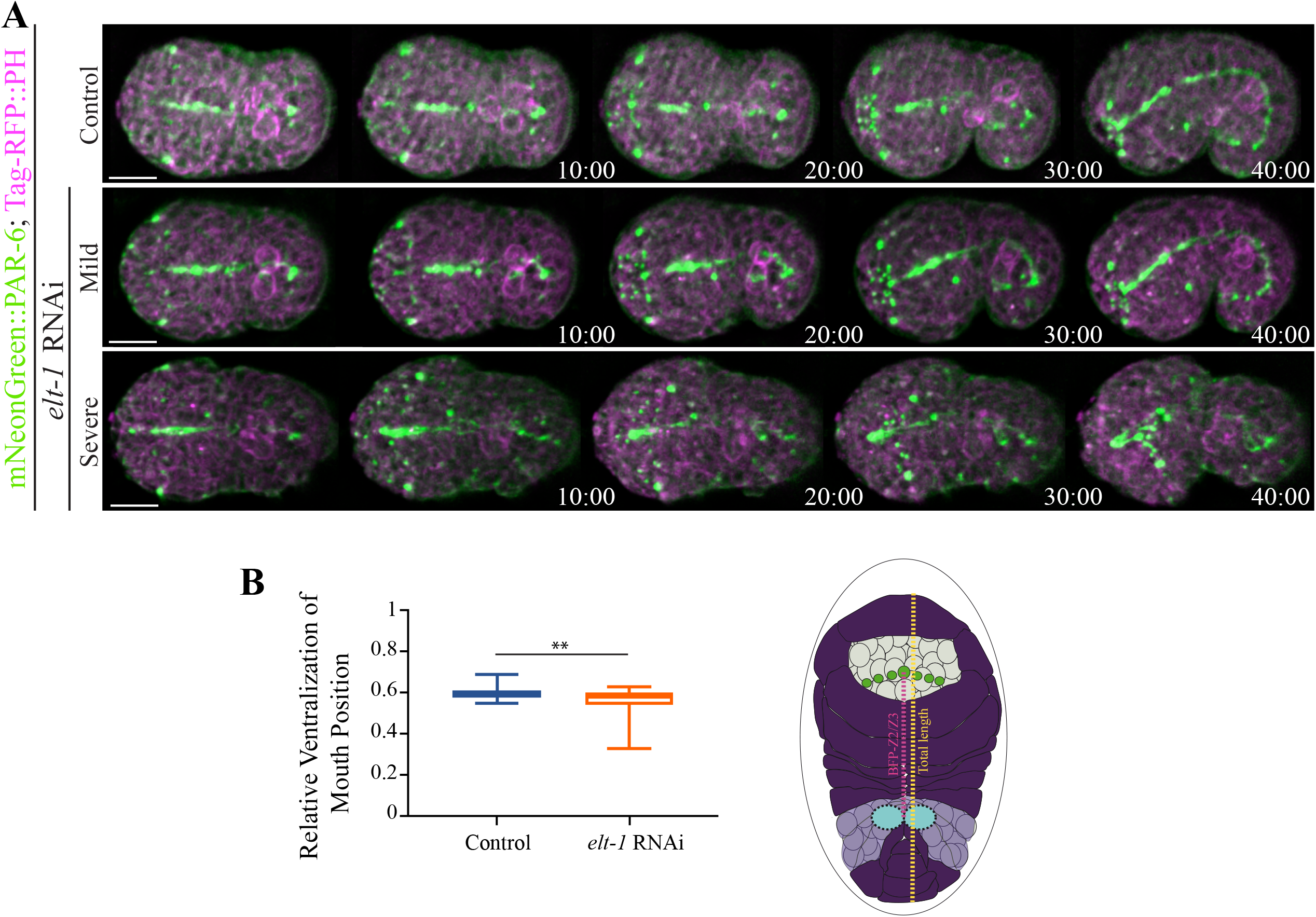
Altering epidermal cell fate causes a change in lumen position. **A)** Time-lapse images acquired using sweptfield microscopy show ventral views (anterior to the left) of embryos expressing mNeonGreen∷PAR-6 (green); Tag-RFP∷PH (magenta) in control (top panel) and *elt-1* RNAi embryos (middle and bottom panels; time in minutes). The scale bar is 10 μm. A range of phenotypes was observed: 1) mild, in which PAR-6+ foci patterns were similar to those of control embryos, although there was a shift toward a more ventrally positioned lumen, and 2) severe, in which foci failed to coalesce into patterns, and the lumen was extremely displaced. **B)** A graph shows the average ratio of the distance between the lumen and Z2/Z3 cells (magenta in the cartoon schematic; future germline) to the total length of the embryo (yellow in the cartoon schematic). Only mildly perturbed embryos were measured, and the variability of the ratio was significantly increased in *elt-1* RNAi (orange) as compared with control (blue) embryos. ** *p* < 0.01.

## Discussion

Here we describe anterior morphogenesis of the *C. elegans* embryo. This developmental process is crucial for giving rise to a properly formed anterior lumen, which must align precisely with the lumen of the pharynx and intestine. Cells from three different tissues are coordinated for anterior morphogenesis: the neuronal cells, pharyngeal cells and epidermal cells. But how they do this is not known, largely due to the complexity of studying multiple tissues simultaneously *in vivo*. We used different types of microscopy and post-acquisition software to define the patterns formed by different cell types during anterior morphogenesis. We observed the ingression of neuroblasts as epidermal cells migrated anteriorly to enclose the anterior surface of the embryo in a layer of epidermis. The neuroblasts were arranged into a circle-like pattern that likely facilitates their inward movement. We propose that their movement could in part be due to anterior-ward forces generated during ventral enclosure. During this stage of epidermal morphogenesis, the ventral surface of the embryo is enclosed by a layer of epidermal cells. We also observed that the arcade cells, the anterior-most pharyngeal cells, formed a rosette after they moved toward the anterior surface and then back into the embryo to a depth of ~5 μm. They formed projections near the anterior region, and we propose that these are the first markers of the location of the future lumen. The ends of the projections accumulated components of apical polarity including PAR-6, NMY-2 and HMP-1. Given that they coalesce, we call them the ‘bright focal point’ (BFP), with the understanding that some signals could be associated with nearby neuronal or pharyngeal cells. Soon after the appearance of the BFP, PAR-6+ve, HMP-1+ve, NMY-2+ve and ANI-1+ve foci, most likely reflecting subsets of polarized neuroblasts, arranged themselves into two pentagons that formed on either side of the BFP. Another set of foci formed a semi-circle that corresponded to the projections of ventral epidermal cells as they migrated anteriorly. These PAR-6+ve, HMP-1+ve and NMY-2+ve foci, which also weakly express AJM-1, could be associated with subsets of neuroblasts and/or epidermal cells and reflect points of adhesion. The ventral epidermal cells migrated with, but never crossed, these foci and remained in a more ventral position, whereas the dorsal epidermal cells migrated with the posterior pentagon foci toward the ventrally positioned BFP. We also observed different types of F-actin–rich projections in the migrating epidermal cells, which appeared longer and more dynamic in the ventral cells than in the dorsal cells, where they were shorter and more uniform in appearance. We found that the neuroblasts were required for controlling the speed of epidermal migration, as reducing their numbers by *ani-1* RNAi caused varying delays. In contrast, mildly perturbing epidermal cell fate caused a shift in the location of the future lumen but did not affect the timing of its formation.

It is not clear how the neuroblasts could affect epidermal cell migration. As described earlier, signaling pathways that regulate WSP-1 and WVE-1 to control the formation of branched F-actin have been implicated in ventral epidermal cell migration during ventral enclosure, and it is likely that they could influence their anterior migration as well (Sawa et al., 2003; Withee et al., 2004; Patel et al., 2008; Bernadskaya et al., 2012; Wallace et al., 2018). It is not clear where the ligands for these pathways are located (*e.g.*, Bernadskaya et al., 2012), but we favor the hypothesis that they originate from the subset of neuroblasts that form the pentagon and/or semi-circle around the BFP. These pathways include those responding to ligands such as UNC-6 (netrin), SLT-1 (slit), EFN-1–4 (ephrins) and MAB-20 (semaphorin) and their receptors UNC-40 (DCC), SAX-3 (Robo), VAB-1 (Eph receptor) and PLX-2 (plexin), respectively (Chin-Sang et al., 1999; George et al., 1998; Wang et al., 1999; Chin-Sang and Chisholm, 2000; Roy et al., 2000; Bernadskaya et al., 2012; Ikegami et al., 2012). We are testing this by perturbing components of these different pathways to determine their impact on anterior epidermal cell migration. As the dorsal cells migrate over a longer distance, they could be influenced by some of the same pathways but with different sensitivities and/or subsets of receptors to account for this distance.

We also propose that there are cues associated with the projections from the arcade cells and/or surrounding neuroblasts that form the BFP. While PAR-6 localization to the BFP was not altered by *ani-1* RNAi, it was decreased by *pha-4* RNAi to change the fate of pharyngeal cells (Kalb et al., 1998; Fig. S1B). Since the BFP is visible before the patterning of neuroblasts, cues associated with these projections could influence neuroblast patterning. While we see ventral shifts in the position of the BFP when we perturb epidermal cell fate, this only occurs at a later timepoint, during polarization of the anterior pharynx. Prior to this, it initiates in the correct position. This suggests that the epidermal cells are crucial for reinforcing lumen position, while the neuroblasts control the timing of lumen formation. We also observed shifts in the position of neuroblast foci just prior to polarization, suggesting that they also associate with and rely on the epidermal cells for late vs. early positioning. This was recently described by the Bao lab for amphid neuroblasts, which form rosettes and piggyback on the migrating epidermal cells for their anterior-directed movement (Fan et al., 2019). We will determine whether perturbing pharyngeal cell fate affects epidermal cell migration and/or neuroblast position. We propose that this cluster of projections provides cues for the initial patterning of the neuroblasts, which in turn control the rate of epidermal cell migration.

Our studies show how cells from different tissues are coordinated to give rise to the anterior lumen during development. These findings will facilitate more in-depth studies of how chemical cues organize and coordinate the patterning and movements of the different cell types. Given the challenges of studying tissues in more complex metazoans, our studies provide fundamental knowledge of how different cell types can be coordinated for the development of structures.

## Acknowledgments

We thank Chris Law for help with imaging studies as part of the Centre for Microscopy and Cellular Imaging at Concordia University. We thank Hari Shroff for his generosity and use of the diSPIM at the NIH/NIBIB, and Harshad Vishwasrao at the trans-NIH Advanced Imaging and Microscopy Resource (AIM) for his technical assistance and time. We thank Michael Glotzer (University of Chicago), Jean-Claude Labbe (IRIC), Susan Mango (Biozentrum of the University of Basel) and Richard Roy (McGill University) for reagents. We thank Victoria Richard and Mathew Duguay for their contributions to various aspects of this work. We acknowledge Wyoming INBRE for editing services, a project that is supported in part by a grant from the National Institute of General Medical Sciences (2P20GM103432) from the National Institutes of Health. The content is solely the responsibility of the authors and does not necessarily represent the official views of the National Institutes of Health. This work was funded by a grant from the National Institutes of Health (NIH) to David Fay (R01GM125091). R.C. acknowledges support from the intramural research programs of the National Institute of Biomedical Imaging and Bioengineering, within the National Institutes of Health.

---

**Figure S1. A) The epidermal cells form concentric rings around the lumen.** Time-lapse images acquired using sweptfield microscopy show *en face* views of two different embryos expressing AJM-1∷GFP (Fire LUTs) during anterior morphogenesis (time in minutes). The Fire LUTs depict signal intensity; cool colors, such as blue or violet correspond to lower levels vs. warmer colors such as yellow or white, correspond to higher levels. The scale bar is 10 μm. **B) Disrupting pharyngeal fate causes perturbations in the BFP.** Time-lapse images acquired using sweptfield microscopy show ventral views of embryos expressing mNeonGreen∷PAR-6 (green); Tag-RFP∷PH (magenta) in control (top panel) and after *pha-4* RNAi to disrupt pharyngeal cell fate (bottom panel; time in minutes). The BFP (yellow arrowheads) was less apparent in *pha-4* RNAi embryos compared to control embryos, but not the other PAR-6+ve patterns. The scale bar for all embryos is 10 μm.

**Movie S1.** A time-lapse movie (7 fps) of the embryo from Fig. 1C co-expressing mNeonGreen∷PH to visualize membranes (green) and H2B∷mCherry to visualize nuclei (magenta), imaged by di-SPIM through anterior morphogenesis.

**Movie S2.** A time-lapse movie (7fps) of the embryo from Fig. 4B (top panel) expressing *lin-26p*;LifeAct∷GFP taken using HILO microscopy (anterior down) to see ventral epidermal cell F-actin projections.

**Movie S3.** A time-lapse movie (7 fps) of the embryo from Fig. 4B (bottom panel) expressing *lin-26p*;VAB-10(ABD)∷mCherry taken using HILO microscopy (anterior down) to see dorsal epidermal cell F-actin projections.

