## Supplementary figures and images for "Multi-tissue patterning drives anterior morphogenesis of the *C. elegans* embryo"

### Figure S1

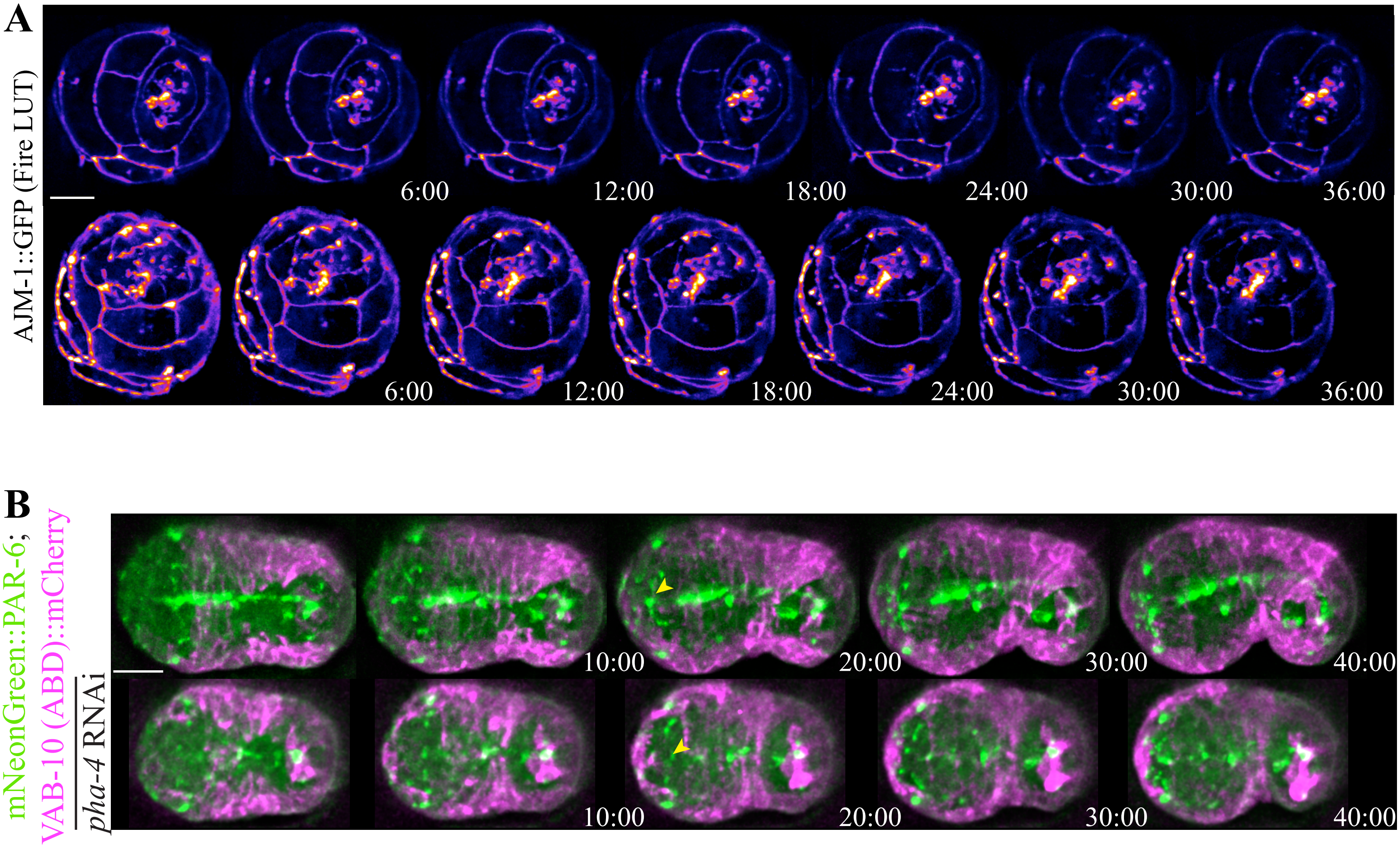
